# Comprehensive analysis of draft genomes of two closely related *Pseudomonas syringae* pv. *syringae* phylogroup 02b strains infecting *Citrus* plants

**DOI:** 10.64898/2025.12.22.696067

**Authors:** Emna Abdellatif, Steve Baeyen, Johan van Vaerenberg, Monika Kałużna, Jaap D. Janse, Ali Rhouma

**Affiliations:** Laboratory of Bioaggressors and Integrated Protection in Agriculture, The National Agronomic Institute of Tunisia, University of Carthage, Tunis, Tunisia; Laboratoire d’Amélioration et Protection des Ressources Génétiques de l’Olivier, Institut de l’Olivier BP 208, Tunis, Tunisia; Flanders Research Institute for Agriculture, Fisheries and Food (ILVO), Plant Sciences, Merelbeke-Melle, Belgium; The National Institute of Horticultural Research, Skierniewice, Poland; Department of Laboratory Methods and Diagnostics, Dutch General Inspection Service (NAK), Emmeloord, Netherlands

**Keywords:** Comparative genomics, Average Nucleotide Identity, secondary metabolites, Type III secretion system (T3SS), copper resistance, prophage analysis, Whole-genome sequencing

## Abstract

*Pseudomonas syringae* pv. *syringae* (*Pss*) is a cosmopolitan phytopathogen with high genomic plasticity and effector diversity, driving host adaptation and variable virulence. We sequenced, assembled, and annotated the genomes of two *Pss* strains, LMG 5496 (from *Citrus sinensis*, Greece, 1962) and EC33 (from *C*. *limon*, Tunisia, 2014), to identify genetic determinants underlying their differential virulence. The genomes of LMG 5496 (6.23 Mb; 58.78% GC) and EC33 (5.98 Mb; 58.97% GC) contained 5,563 and 5,312 coding sequences, respectively, and shared highly conserved metabolic and functional profiles. Average Nucleotide Identity (ANI ≈ 100%) and phylogenomic analyses confirmed their close evolutionary relationship within phylogroup 02b.

Despite overall similarity, EC33 retained a complete repertoire of Type III secretion system (T3SS) effectors, while LMG 5496 exhibited deletions in two core effectors, correlating with reduced virulence. Functional annotation identified 16 high-confidence T3SS effectors, 12 shared and four strain-specific, reflecting divergent molecular strategies. Enhanced pathogenicity of EC33 was linked to the presence of hypervirulence gene Q4ZSY4 and loss of AvrPph3, suggesting host immune evasion. Both strains possessed largely conserved copper resistance genes, with minor differences in detoxification enzymes. Prophage profiling revealed distinct histories: LMG 5496 carried two intact prophages, EC33 one.

This comparative genomic analysis provides new insights into adaptation, phenotypic diversification, and shaping virulence within clonal *P*. *syringae* lineages. To our knowledge, this is the first genome-based comparison of *Pss* strains isolated from *Citrus*.

## Introduction

*Citrus* crops rank among the most valuable fruit commodities worldwide, notably in the Mediterranean basin, including Tunisia, where they underpin both agricultural livelihoods and export revenues (Beshara et al., 2024). Due to the ubiquity of fungal and viral threats in citrus production, bacterial pathogens have historically been under recognized in Tunisian orchards. However, bacterial pathogens are key contributors to the emergence of new plant diseases, not in the last place due to the adaptivity of their genomes under ecological, environmental, and agricultural changes of their habitats (Vos et al., 2024; Maddock and Hulin, 2025).

More recently, outbreaks of the diseases bacterial blast and so-called black pit on lemon and orange have been attributed to *Pseudomonas syringae* pv. *syringae* (*Pss*), highlighting the emergence of a previously overlooked disease complex in this region (Abdellatif et al., 2015, 2017, 2020; Oueslati et al., 2020).

The genus *Pseudomonas*, described by Migula in 1894, encompasses a phylogenetically diverse group of over 300 validly published (Freese et al., 2025) species that thrive across environments and hosts, reflecting their remarkable ecological plasticity (Peix et al., 2018; Yi and Dalpke, 2022; Udaondo et al., 2024). Within this genus, the *P*. *syringae* complex is typified by pathovars (>60) that exploit a broad host range, environmental adaptation and diverse mechanisms of virulence. Central among the latter is the Type III secretion system (T3SS), which injects effector proteins into plant cells to suppress immunity and enable colonization (Xin et al., 2018; Jin et al., 2023; Maguvu et al., 2024).

Given these dynamics, high resolution, genome-based investigations are essential to decipher the determinants of host specificity, virulence, and resistance; particularly in the context of expanding bacterial threats to *Citrus* spp. and other crops and the evolving agro-ecological interfaces in which they operate (Ogbuji and Agogbua, 2025).

In Tunisian *Citrus* systems, molecular and phylogenetic studies have shown that *Pss* strains associated with the bacterial diseases blast and black pit belong to phylogroups PG02b and PG02g; however, they exhibit substantial genetic heterogeneity, highlighting their adaptive potential and complicating disease management (Abdellatif et al., 2020).

To date, the genetic diversity of the *P*. *syringae* population in *Citrus* spp. cultivation remains poorly studied (Abdellatif *et al*., 2017; 2020; Oueslati et al., 2020) although there are several studies devoted to the comparative genomic analysis of *P*. *syringae* pathovars from other crops, such as from tomato, kiwifruit, bean, *Prunus* and pepper (Baltrus et al., 2011; McCann et al., 2013; Nowell et al., 2016; Ruinelli et al., 2019 and 2022; Ranjit et al., 2024).

Strains of *Pseudomonas syringae* (PG 02b) were described to cause bacterial blast and black pit in almost all known citrus-growing regions in Tunisia, based on multilocus sequence analysis (Abdellatif et al., 2020). Two strains, assigned to this clade, were selected for genome analysis: strain EC33, isolated from *Citrus limon* cv. Eurêka in 2014 with symptoms of black pit in the Mornag area near Tunis and shown to be highly virulent in pathogenicity assays and (as reference) strain LMG 5496, isolated in 1962 from *Citrus sinensis* in Greece, showing symptoms of bacterial blast and black pit as well.

The major driver for sequencing the whole-genome of these two strains from *Citrus* was to gain a deeper insight into bacterial function, in terms of key genes responsible for virulence, host specificity prophage content and copper resistance and to compare the genomes of these strains (differing in host, isolation place and time) with other available genomes of PG02, based on the calculation of the average nucleotide identity (ANI).

## Materials and Methods

### Strains and pathogenicity tests

Strain EC33 was isolated from *Citrus limon* (cv. Eurêka) at the Laboratory of Bioaggressors and Integrated Pest Management in Agriculture in the National Agronomic Institute of Tunis-Tunisia in 2014 (Abdellatif et al., 2017). The strain was highly virulent and representative of a Tunisian *Pss Citrus* collection of 2014-2015. Strain LMG 5496 was isolated from *C. sinensis* by P. Panogopoulous in Greece in 1962 (Panagopoulos, 1964). Both strains were stored at - 80°C in sterile distilled water with 15% glycerol. Before use, they were cultured onto King’s B agar medium for 48 h at 28°C. Their phenotypic, genotypic and pathogenicity characterization was performed in a previous study (Abdellatif et al., 2017; 2020).

### DNA extraction and genome sequencing, assembly, and annotation

Genomic DNA was isolated in the Laboratory of Bacteriology in the Flanders Research Institute for Agriculture, Fisheries and Food Plant Sciences Unit, Belgium from the strains via the NucleoSpin® Microbial DNA kit. The DNA was shipped to BGI, Hong Kong, where the Illumina HiSeq 4000 platform was used to generate 100-bp paired end reads. A total of over 1GB sequence data was obtained for both strains in 300 bp insert-size PCR-free libraries. De novo assembly with SeqMan Ngen v.14.1.0.115 (DNASTAR) was done, followed by decontamination of the draft genome contigs with Blobtools version I (Laetsch and Baxter, 2017).

### Data availability

The genome sequences for EC33 and LMG 5496 are available under NCBI BioProject number PRJNA723174, with annotated assemblies available under accession numbers JAGSOX000000000 and JAGSOW000000000, respectively. The Sequence Read Archive (SRA) accession numbers are listed in Table 1. Sequence reads were deposited in the NCBI SRA under the accession numbers SRR14286381 and SRR14286382.

**Table 1.**
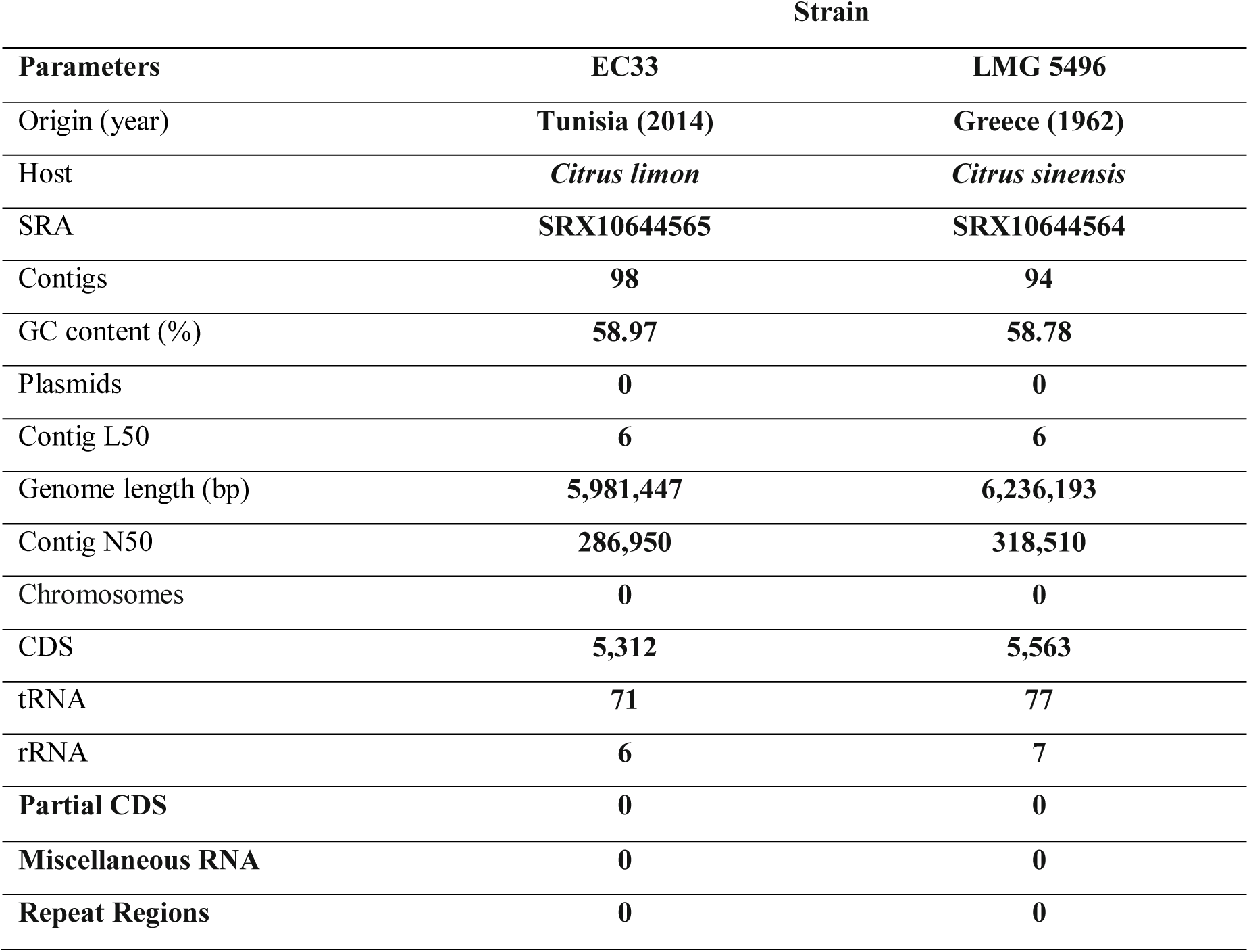
Key parameters of the assembly and annotation of the genomes of *Pseudomonas syringae* pv. *Syringae* strains EC33 and LMG5496.

### Comparative Genomics and Phylogenetic Analysis

#### 1. Genome assembly and annotation

The genome assemblies of *Pseudomonas syringae* strains LMG 5496 and EC33 were analysed using the Comprehensive Genome Analysis service of the Bacterial and Viral Bioinformatics Resource Centre (BV-BRC v3.54.6, https://www.bv-brc.org/, Wattam et al., 2017). Following submission to BV-BRC, the genomes were annotated and quality-checked through the PATRIC pipeline. According to the annotation statistics and comparison with other *P*. *syringae* genomes available in BV-BRC, both assemblies exhibited high completeness and overall good quality.

The *Pss* EC33 and LMG 5496 genomes were annotated using the RAST tool kit (RASTtk) and the genetic code 11 (Brettin et al., 2015) and assigned unique genome identifiers of 321.326 and 321.325, respectively.

#### 2. Average Nucleotide Identity (ANI) Analysis

Average Nucleotide Identity (ANI) analysis was performed to assess the genomic similarity among 100 *Pseudomonas* strains, including the two newly sequenced strains from *Citrus*, EC33 and LMG 5496. Pairwise genome similarity was evaluated using the ANIm algorithm implemented in pyANI-plus v1.0.0 (https://pyani-plus.github.io/pyani-plus-docs/). The draft genomes were compared against reference genomes of closely related *Pseudomonas* species and pathovars within *Pseudomonas syringae* obtained from NCBI RefSeq entries in BV-BRC, using their command-line interface CLI v1.048. The Overall genome relatedness index (OGRI) method ANIm was used to calculate the values based on MUMmer (NUCmer) whole-genome alignments, with coverage and identity matrices extracted for clustering and visualization. A species boundary threshold of 95% ANI was applied to delineate species-level relationships.

A percentage identity heatmap was generated from the pairwise ANIm values (available as tsv file) using the pyani-plus subcommand anim and the host metadata using the R script create_ani_complexheatmap.R.

#### 2. Effector Data Retrieval and Analysis

A total of 161 effector-associated genes, linked to *Pseudomonas* species and pathovars within *Pseudomonas syringae* were retrieved from the Pathogen-Host Interactions database (PHI-base, v4.18; www.phi-base.org). Relevant effectors for *Pseudomonas* were identified by querying the PHI database with the organism’s taxonomic identifier and filtering for genes annotated with effector or virulence-associated phenotypes using the script fasta_filter.py. Protein sequences of identified effectors were subjected to tblastn searches against the genomes using the Blast+ v2.17.0 suite with E-value threshold of 1e-5 and minimum of 60% coverage as cut-offs using the Python script blast_matrix_script.py to generate a binary matrix file PHI_binary_matrix.csv. Effector families were categorized based on functional domains and known virulence mechanisms.

Effector genes were classified according to PHI-base phenotype annotations: (i) reduced virulence, (ii) loss of pathogenicity, (iii) avirulence effector, (iv) increased virulence, (v) unaffected pathogenicity, and (vi) mixed effect. The binary matrix file in csv format was finally used to generate a hierarchically clustered heatmap in RStudio v2025.09.0+387 (https://dailies.rstudio.com/version/2025.09.0+387/) with R v4.5.0 and the package Complex Heatmap v2.24.1, using the script PHI_heatmap_with_host.R. For multi-strain (100 strains) comparisons, presence or absence of effectors across phenotypes was encoded numerically (0 = absent, 1 = present). Hierarchical clustering was applied to both rows (effectors) and columns (phenotypes/strains) using Euclidean distance and complete linkage to reveal patterns of similarity. Color gradients were selected to distinguish phenotypic categories. Row and column dendrograms were displayed to facilitate the identification of effector clusters with similar functional profiles. Specific annotations, such as effector families or host specificity, were added as sidebars.

#### 3. Type III Effectors Prediction (Effectidor)

Protein-coding sequences (CDSs) from the draft genomes of *Pss* strain EC33 and strain LMG 5496 were analyzed using Effectidor II (https://effectidor.tau.ac.il, Wagner et al., 2025), a pan-genomic AI-based algorithm for the prediction of Type III secretion system (T3SS) effectors. The tool integrates sequence features, structural motifs, and homology to known T3Es, to assign a probability score to each protein. The Effectidor II platform was used to predict potential T3SS effectors in the bacterial genomes under study. Predictions with a confidence score ≥ 0.90 were considered high-confidence effector candidates. Functional annotations and effector family assignments were derived from the closest homologs in the genus *Pseudomonas* and related plant-pathogenic bacteria. Redundant or truncated predictions were removed based on sequence overlap and manual curation of protein function.

#### 4. Biosynthetic gene cluster prediction (antiSMASH Analysis)

Prediction of secondary-metabolite biosynthetic gene clusters (BGCs) was performed using antiSMASH version 8.0.4 (https://antismash.secondarymetabolites.org). Analyses were conducted using relaxed detection parameters to maximize identification of potential BGCs. The KnownClusterBlast function was enabled to allow comparison with previously characterized clusters in the MIBiG database. Default settings were applied for cluster border prediction, domain annotation, and functional assignment. Identified BGCs were categorized according to predicted biosynthetic type, including NRPS, PKS, hybrid NRPS–PKS, trans-AT PKS, siderophore/NRP-metallophore, arylpolyene, bacteriocin, terpene, and cryptic NRPS-like clusters. Cluster location (contig and coordinates), predicted product type, and similarity to known clusters were extracted from the antiSMASH output.

#### 5. Copper Resistance Genes (metHMMDB Analysis)

Copper resistance genes were identified by querying protein sequences of the two studied *Pseudomona*s strains against the MetHMMDB database (https://methmmdb.com/, Ciuchciński and Dziurzyński, 2025), a curated repository of 254 profile Hidden Markov Models (HMM’s) for microbial metal resistance genes. Searches were performed using HMMER (v3.4) and the 52 HMMER models from MetHMMDB related to Cu copper resistance. In short, the script run_hmmscan_parallel.sh was used to search the CDS of the genomes in amino acid format with an E-value threshold of=1e-5. The HMMER output files were parsed with the script parse_hmm_results.sh, that uses the python script HmmPy with an E-value threshold 1e-5 and a minimum coverage of 40%. A binary matrix file (hmmscan_binary_matrix.tsv) was then generated, using the Python script hmm_results_to_binary_matrix.py, to use as input in order to generate a hierarchically clustered heatmap with the R-script copper_resistance_heatmap-with_host.R. Identified genes were annotated according to functional domains and categorized into known copper resistance mechanisms, including Cu-efflux pumps (*CopA*, *CopB*), multicopper oxidases (*CueO*), and copper chaperones. Results were compiled in tabular format using R v4.5.0 and the ComplexHeatmap module for further comparative analysis across the 100 strains.

This allowed the identification of key copper resistance determinants, including multicopper oxidases, copper-exporting P-type ATPases, and RND efflux components (e.g., CusCBA).

#### 6. Prophage identification and classification

Prophage regions were identified using the PHASTEST web server (Wishart et al., 2023; https://phastest.ca). The complete genome assemblies of strains LMG 5496 and EC33 were uploaded to the PHASTEST submission form in FASTA format and analyzed using the server default settings (accessed 18 September 2025). Prophage prediction was performed using default PHASTEST parameters, which combine sequence similarity searches and structural gene detection to delineate putative prophage regions. Each detected region was automatically assigned a completeness score (0–150) and classified according to PHASTEST criteria as: intact (score ≥ 90), questionable (score 70–90), or incomplete (score < 70) (as described in Wishart et al., 2023).

For each identified prophage region, the following parameters were extracted from the PHASTEST output: genomic coordinates (start–end positions), prophage length (bp), GC content (%), number of predicted proteins, closest phage reference (based on best BLAST hit), and PHASTEST completeness score. The prophage repertoires of the two *Pss* strains LMG5496 and EC33 were then compared based on number, size, gene content, and phage identity. Comparative interpretation focused on differences in prophage composition as indicators of strain-specific infection history, prophage acquisition, retention or loss, and genomic evolution following divergence.

## Data availability

Following the FAIR principles, all data including genome files in nucleotide and amino acid format, scripts and result files for the PHI-base, pyani-plus and MetHMMDB analyses are available at https://gitlab.ilvo.be/stevebaeyen/pseudomonas_genomes_citrus_2025.

## Results

### Sequencing, de novo assembly, and annotation of strains

Genome analyses showed that LMG 5496 was assembled into 94 contigs (total length 6,236,193 bp; GC content 58.78%), whereas EC33 comprised 98 contigs (5,981,447 bp; GC content 58.97%) (Table 1). Both genomes displayed similar assembly quality, with L50 (number of contigs covering 50% of the genome) values of 6 and N50 (length of the contig that covers 50% of the genome) values of 318,510 bp (LMG 5496) and 286,950 bp (EC33). No plasmids were detected in either isolate. RASTtk-based annotation predicted 5,563 CDS, 77 tRNA, and 7 rRNA genes in LMG 5496, while for EC33 the prediction was 5,312 CDS, 71 tRNA, and 6 rRNA genes (Table 1).

Annotation using RASTtk predicted 5,563 coding sequences (CDS), 77 tRNA, and 7 rRNA genes in LMG 5496, and 5,312 CDS, 71 tRNA, and 6 rRNA genes in EC33 (Table 1). LMG5496 contained 1,106 hypothetical proteins, while EC33 harbored 1,268 of those, indicating a modest increase in uncharacterized genes in EC33. The number of functionally annotated proteins was comparable between strains (4,206 in LMG 5496 vs. 4,295 in EC33). Minor differences were also noted in functional assignments (EC-number, GO, and pathway annotations), likely reflecting strain-specific metabolic differences rather than annotation bias (Table 2 and Figure 1). Subsystem analysis revealed similar overall functional profiles. Metabolism-related genes dominated both genomes (969 in LMG 5496 and 951 in EC33), followed by protein processing, stress response, virulence, membrane transport, and energy metabolism categories (Figure 2), indicating broadly conserved physiological capacities.

**Figure 1.**
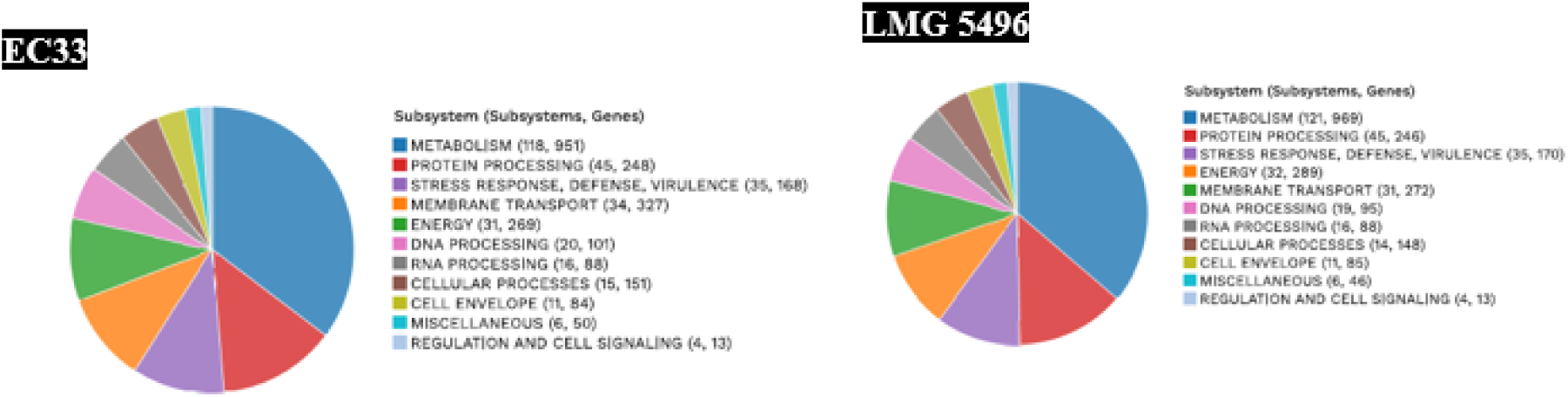
Schematic overview of subsystem coverage, subsystem category distribution, and subsystem feature counts in the annotated genome of *P*. *syringae* pv. *syringae* strains LMG 5496 and EC33. The prediction used SEED Viewer v2.0. The graphic was generated using RAST SEED Viewer v.2.0. Genomic features are color-coded according to their functional classification.

**Figure 2.**
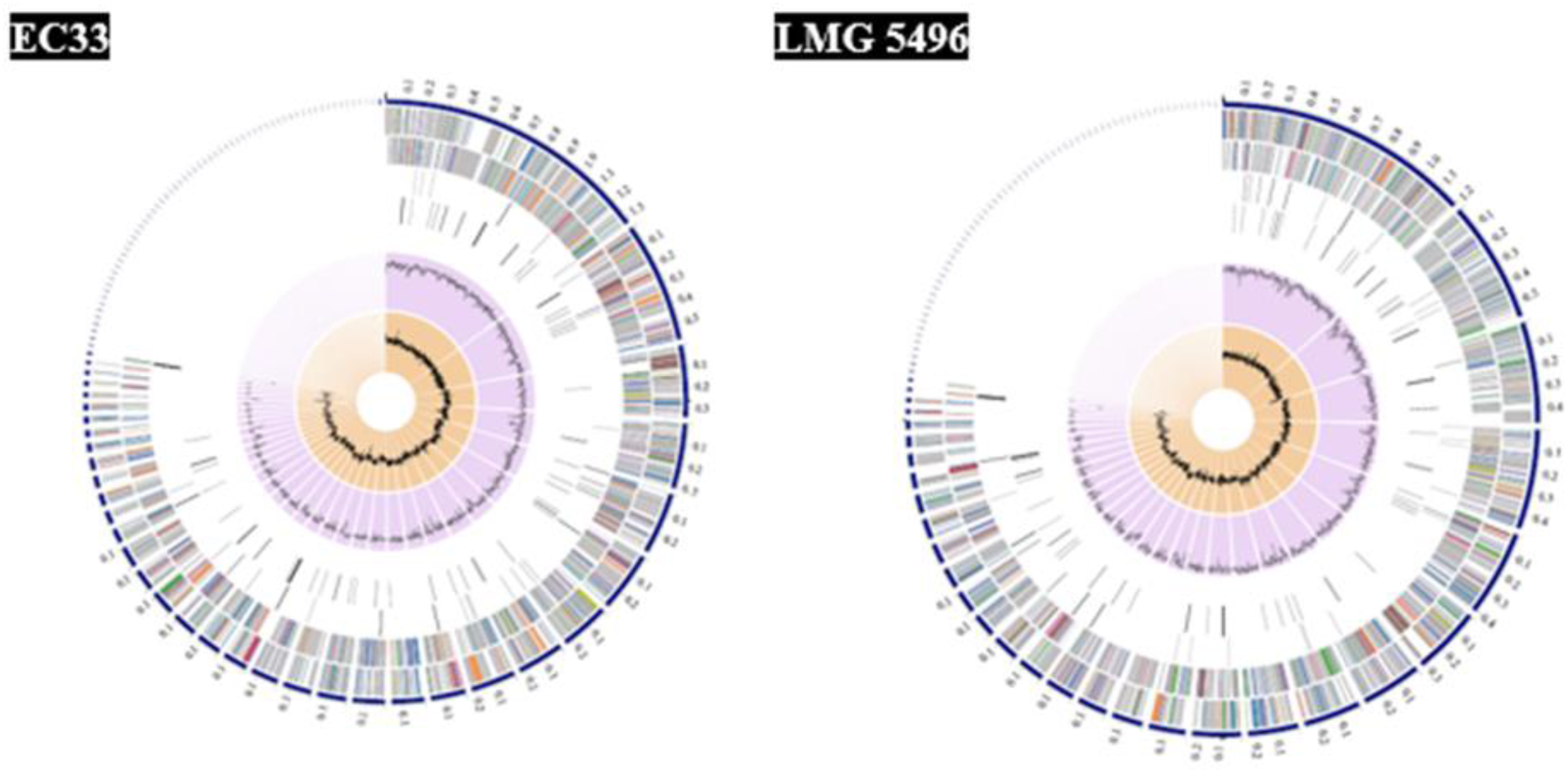
Circular visualization of bacterial genome annotations. The genome map was generated using the Bacterial and Viral Bioinformatics Resource Center (BV-BRC; https://www.bv-brc.org/). From the outermost to the innermost rings, the diagram displays assembled contigs, coding sequences (CDSs) on the forward and reverse strands, RNA-encoding genes, CDSs associated with antimicrobial resistance genes, CDSs associated with virulence factors, GC content, and GC skew. Coding sequences are color-coded according to their assigned functional subsystems.

**Table 2:** Protein Features.

| Protein Features | EC33 | LMG 5496 |
| --- | --- | --- |
| **Hypothetical proteins** | **1,106** | **1,268** |
| **Proteins with functional assignments** | **4,206** | **4,295** |
| **Proteins with EC number assignments** | **1,149** | **1,174** |
| **Proteins with GO assignments** | **976** | **1,001** |
| **Proteins with Pathway assignments** | **857** | **880** |
| **Proteins with PATRIC genus-specific family (PLfam) assignments** | **5,093** | **5,297** |
| **Proteins with PATRIC cross-genus family (PGfam) assignments** | **5,153** | **5,378** |

### Pairwise Average Nucleotide Identity

Pairwise ANI comparisons indicated a high genomic similarity (≈ 100%) between the two isolates, confirming their classification as *P*. *syringae* pv. *syringae*. Phylogenetic reconstruction using concatenated PGFam alignments and RAxML clustered LMG 5496 and EC33 with reference *Pss* strains, supporting their evolutionary relatedness (Figure 3). The ANI value close to 100% between LMG 5496 and EC33 suggests they represent the same clonal lineage or highly conserved variants within it.

**Figure 3.**
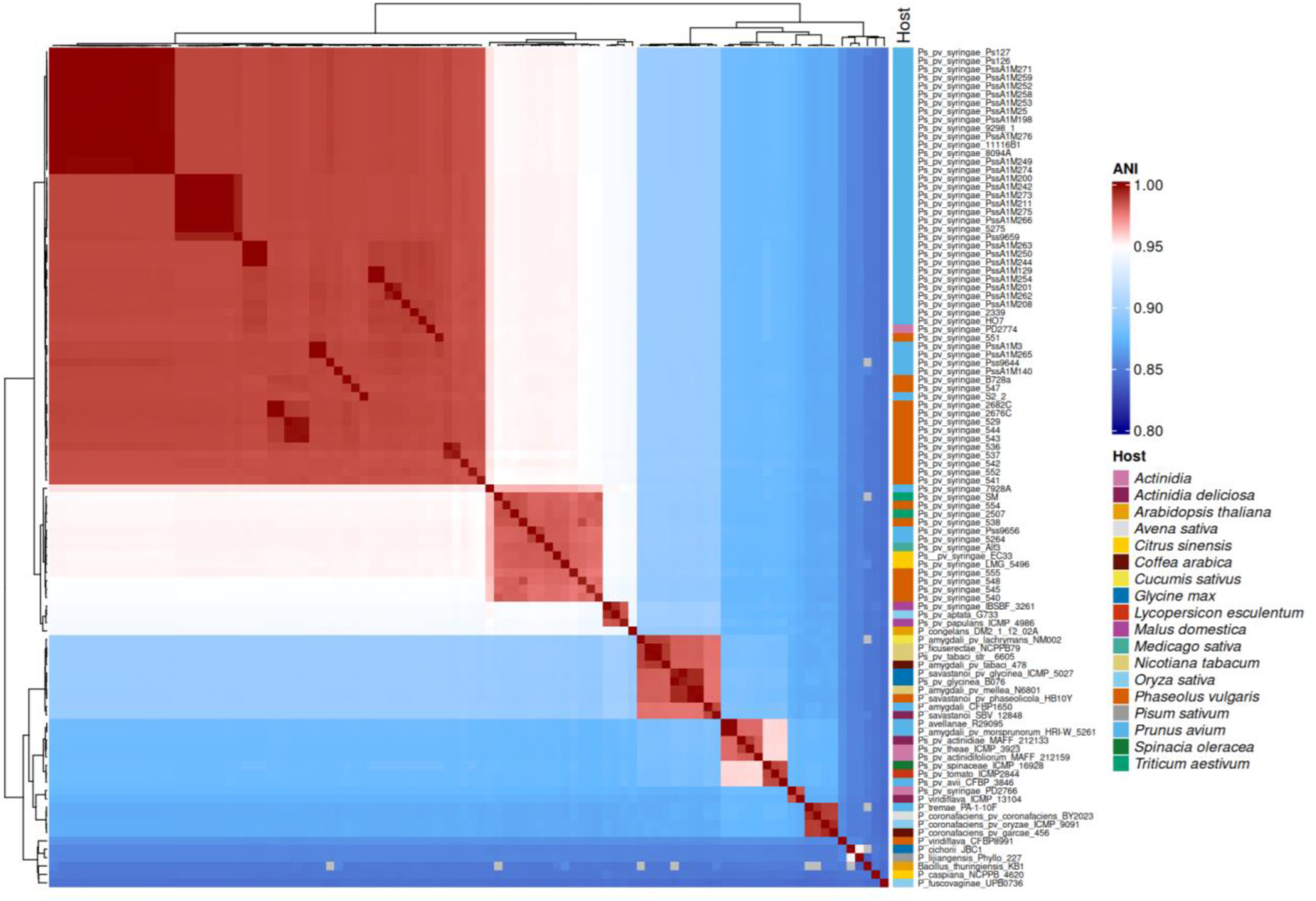
Heatmap of pairwise Average Nucleotide Identity (ANIm) values among *Pseudomonas* strains. The heatmap illustrates the genome similarity matrix calculated using the ANIm algorithm implemented in pyANI-plus v1.0.0.

### Specialty Genes, Virulence, and Antimicrobial Resistance

Both genomes harbored genes associated with transport, virulence, and antibiotic resistance. LMG 5496 encoded 33 virulence factors, while EC33 carried 34; according to VFDB database. Antimicrobial resistance (AMR) genes were largely shared, including efflux pumps (e.g., *MexAB*-*OprM*, *EmrAB*-*TolC*), antibiotic target modification or replacement genes, and oxidative stress regulators such as *OxyR* (Table 3). Analysis across multiple databases revealed subtle differences in AMR potential: both strains carried 1 gene in NDARO, 6 in CARD, and 66–67 in PATRIC, with EC33 slightly lower than LMG 5496 in PATRIC (Table 4). Additionally, the two genomes exhibited comparable profiles for drug targets, metal resistance, and transporters, with 14–15 drug target genes (TTD), 7 metal resistance genes (BacMet), and 78–79 transporters (TCDB).

**Table 3:**
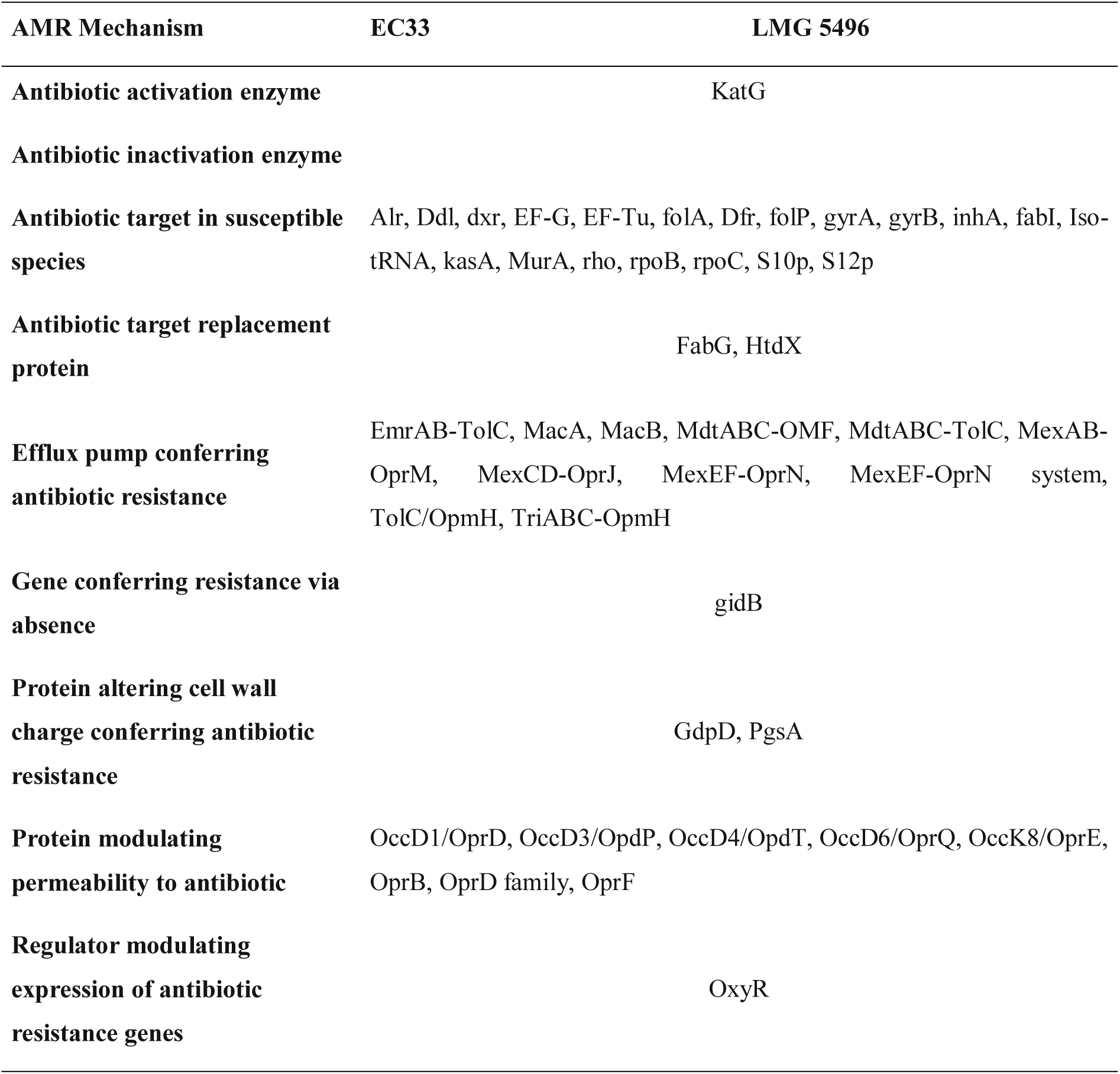
Antimicrobial Resistance Genes.

**Table 4.** Specialty Genes in two *Pseudomonas syringae* pv. *syringae* strains EC33 and LMG5496.

|  | Source | EC33 | LMG 5496 |
| --- | --- | --- | --- |
|  | NDARO | 1 | 1 |
| Antibiotic Resistance | CARD | 6 | 6 |
| Antibiotic Resistance | PATRIC | 66 | 67 |
| Drug Target | TTD | 15 | 14 |
| Metal Resistance | BacMet | 7 | 7 |
| Transporter | TCDB | 78 | 79 |
| Virulence factor | VFDB | 34 | 33 |

### Micro-Evolutionary Divergence in a Clonal Lineage

Despite their nearly identical ANI value of (≈ 100%, a notable divergence was observed within the conserved part of the genome where virulence factors are located. Strain LMG 5496 lacked specific genes present in EC33 and most other *P*. *syringae* strains (Figure 4). These missing genes are located within the Type III Secretion System (T3SS) pathogenicity island.

**Figure 4.**
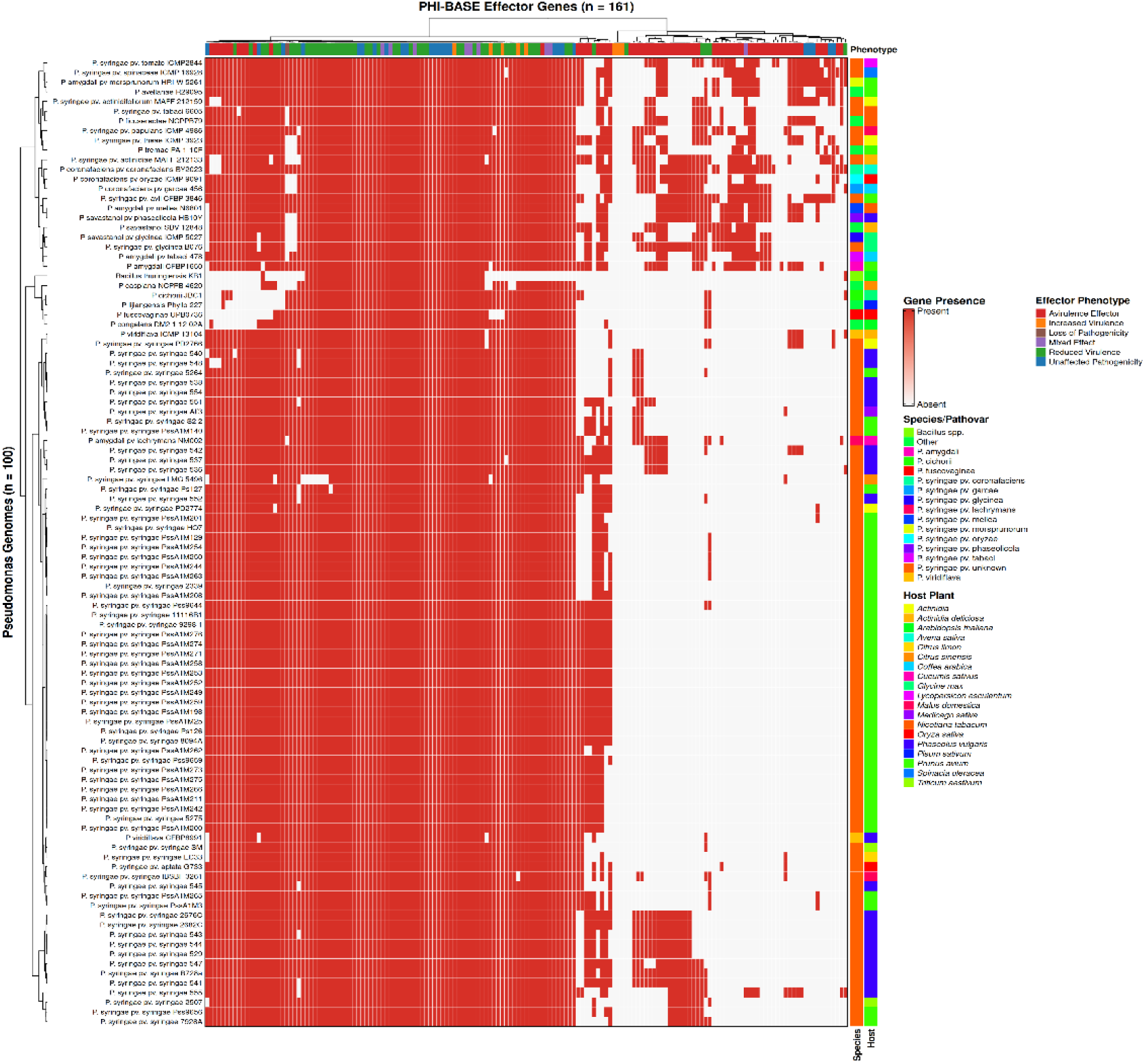
Heatmap of PHI-base Effector Distribution in *Pseudomonas* spp. The heatmap represents 161 effector genes retrieved from PHI-base (v4.18), clustered according to their association with experimentally validated phenotypic categories: reduced virulence, loss of pathogenicity, increased virulence, effector, unaffected pathogenicity, and mixed effect. Rows correspond to *Pseudomonas* pathovars, and columns correspond to PHI-base effector genes. Hierarchical clustering was performed using RStudio v2025.09.0+387 with R v4.5.0 and the package Complex Heatmap v2.24.1. Color intensity indicates the presence / absence of each effector.

Among 161 *Pseudomonas syringae* effector-associated genes identified in PHI-base, approximately 37.9 % corresponded to core T3SS components, 59.0% to variable host-specific effectors, and around 3.1% to unique strain-specific ones. Both genomes retained the core hrp/hrc cluster (e.g., *HrpR*, *HrpS*, *HrpG*, *AlgU*, *GacA*, *Vfr*), confirming an intact T3SS machinery.

Heatmap clustering of virulence factor sequences revealed a clear divergence: EC33 possessed a complete set of core and accessory T3SS effectors, while LMG 5496 lacked two highly conserved core effectors, see below. This microevolutionary loss correlates with phenotypic data (Abdellatif et al., 2017), where EC33 exhibited higher virulence than the effector-deficient LMG 5496. Specifically, EC33 retained ten key virulence genes, including the hypervirulence determinant Q4ZSY4 (*Bsi*), and lost the avirulence gene Q52430 (*AvrPph3*). Conversely, LMG 5496’s reduced virulence likely results from the loss of Q4ZSY4, while retaining Q52430 (triggering host recognition) and Q83Y51 (*SylA*) (Table 5). These findings suggest that EC33’s increased virulence derives from adaptive gene loss enhancing both immune evasion and metabolic efficiency.

**Table 5:** Comparative analysis and functional significance of virulence gene divergence in two *Pseudomonas syringae pv. syringae* strains LMG5496 and EC33, as annotated using the PHI-base.

| Effector Gene (ID #PHI) | PHI-BASE Annotation | EC33 Status (High Virulence) | LMG 5496 Status (Reduced Virulence) | Functional Significance and Evolutionary Impact |  |
| --- | --- | --- | --- | --- | --- |
| Q4ZSY4 (Bsi) | Increased virulence (hypervirulence) | Present (1) | Absent (0) | **Driver of EC33's high pathogenicity:** The presence of this hypervirulence gene is the main molecular determinant for EC33's high pathogenicity. Its loss is the primary reason for LMG 5496's reduced virulence. | McGrath et al. |
| Q7PC62 (HopAE1) | Effector (plant avirulence determinant) | Present (1) | Absent (0) | **Reduced Pathogenicity:** This gene is generally required for full pathogenic fitness; its absence contributes to a less effective infection pathway in LMG 5496. | Vinatzos et al. |
| Q87UQ6 (PSPTO_5240) | Unaffected pathogenicity | Present (1) | Absent (0) | Likely a neutral loss in LMG 5496, but its presence is part of EC33's overall conserved repertoire. | Lam et al. |
| 7 Genes (Q87VY6 to Q88AM7) | Reduced virulence | Present (1) | Absent (0) | **Cumulative Fitness Loss:** These genes are generally required for full pathogenic fitness; their cumulative loss in LMG 5496 leads to a less effective infection strategy. | Liu et al. & Turner et al. |
| Q52430 (AvrPph3) | Effector (plant avirulence determinant) | Absent (0) | Present (1) | **EC33's Evasion Strategy:** EC33 strategically shed this gene to remove a host recognition flag, potentially broadening its effective host range. LMG 5496's retention makes it vulnerable to host R-genes. | Simoni et al. |
| Q83Y51 (SylA) | Reduced virulence | Absent (0) | Present (1) | **EC33's Efficiency Strategy:** This gene is involved in syringomycin toxin synthesis. EC33 | Groll et al. |

| Effector Gene (ID #PHI) | PHI-BASE Annotation | EC33 Status (High Virulence) | LMG 5496 Status (Reduced Virulence) | Functional Significance and Evolutionary Impact |
| --- | --- | --- | --- | --- |
|  |  |  |  | shed it to reduce a high metabolic cost or an evolutionary burden. LMG 5496's retention reflects an older, less optimized state. |

**Table 6:**
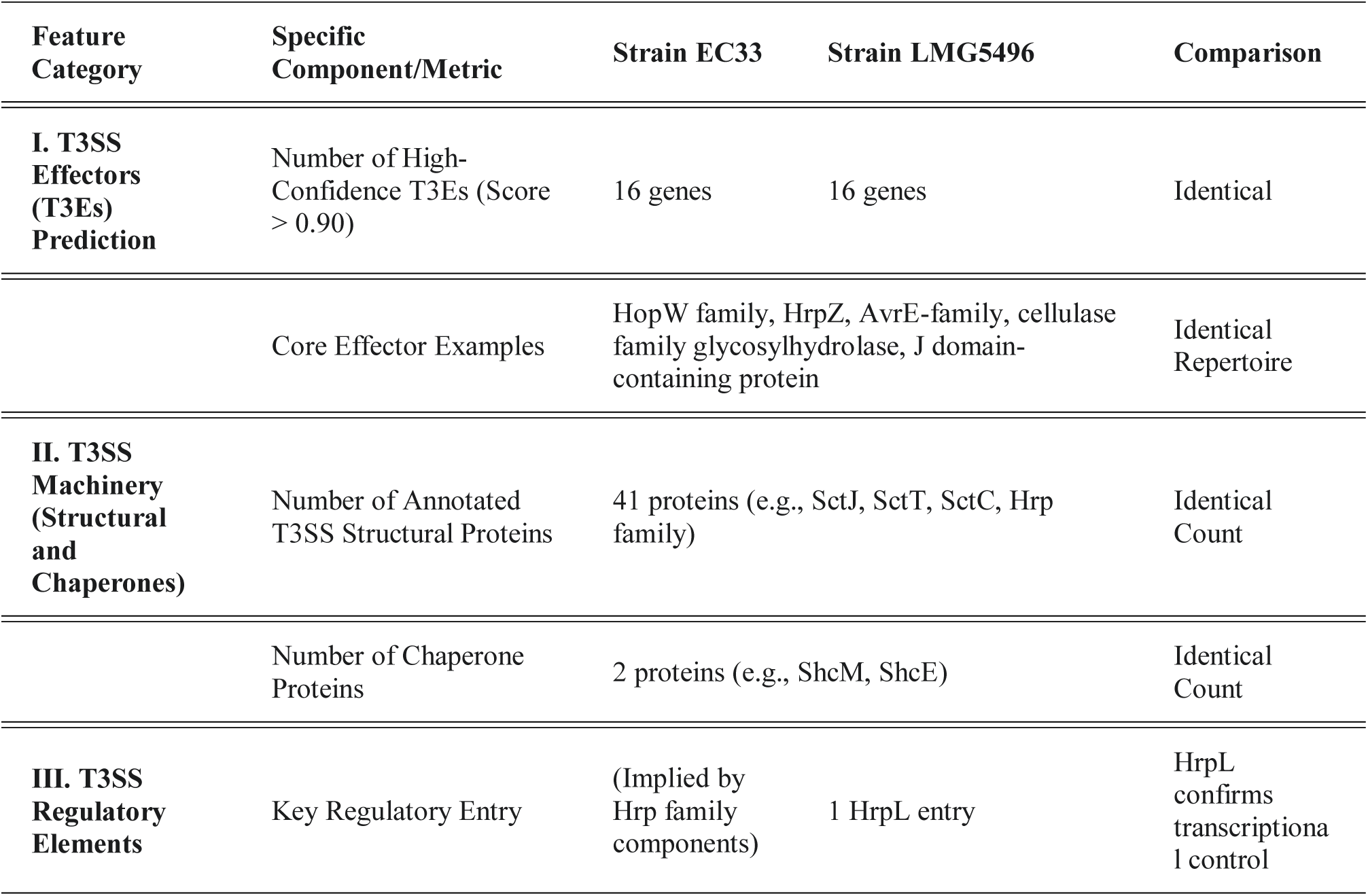
Assessment of Functional Type III Secretion System (T3SS) for two *Pseudomonas syringae pv. Syringae* strains LMG5496 and EC33.

### Type III Effectors Prediction (using Effectidor II)

Effectidor II analysis of *Pss* strains LMG 5496 and EC33 predicted 16 high-confidence Type III effectors in each strain (score ≥0.9), including canonical effectors HopW, HrpZ, AvrE, HopA1, HopAK1, and accessory proteins such as cellulase, pectate lyase, and membrane-targeted toxins. Two chaperone proteins (ShcM and ShcE) and 41 T3SS structural and regulatory components were identified in both strains, supporting the presence of a functional Type III Secretion System. All high-confidence effectors were shared between EC33 and LMG 5496, with no strain-specific differences observed in this analysis (Table6).

### Comparative overview of secondary-metabolite biosynthetic gene clusters (BGCs)

AntiSMASH analysis identified 15 putative secondary-metabolite regions in EC33 and 18 regions in LMG 5496. Both genomes encode NRPS clusters, an NRPS–PKS (syringolin-like) hybrid and a trans-AT PKS (secimide-like), siderophore/NRP-metallophore loci and arylpolyene regions; LMG 5496 additionally contains a larger set of distinct NRPS/NRPS-like clusters (including nunapeptin/nunamycin- and kolossin-like hits). Table 7 summarizes all biosynthetic gene cluster (BGC) classes detected in the two strains using antiSMASH.

**Table 7:** Comparative overview of secondary-metabolite biosynthetic gene clusters (BGCs) identified in *Pseudomonas syringae* pv. *syringae* strains EC33 and LMG 5496 using antiSMASH.

| BGC class | EC33 | LMG 5496 | Notes |
| --- | --- | --- | --- |
| **NRPS (Type I)** | Present. Regions with high similarity to syringomycin, rhizomide, bovienimide (e.g. Region 1.1, 31.1, 12.1) | Present. Multiple NRPS regions with matches to syringomycin, kolossin, ririwpeptide, bovienimide (e.g. Region 2.1, 12.1, 29.1, 32.1) | Both strains carry several NRPS clusters; LMG 5496 shows more distinct NRPS hits across regions. |
| **NRPS-PKS hybrid (NRPS + T1-PKS / NRPS+PKS)** | Present — e.g. Region 6.1 with strong similarity to syringolin A (NRPS:Type I + PKS) | Present — e.g. Region 1.1 with high similarity to syringolin A (NRPS:Type I + PKS) | Both genomes carry a syringolin-like hybrid cluster (conserved match). |
| **trans-AT PKS (PKS:Trans-AT)** | Present — Region 2.1 matches secimide (high confidence) | Present — Region 5.1 matches secimide (high confidence) | Both strains have trans-AT PKS clusters with secimide-like similarity. |
| **Siderophore / NRP-metallophore** | Present — NI-siderophore + NRPS region (Region 1.1 listed as NI-siderophore/NRPS); Region 14.1 flagged as NRP-metallophore (low confidence, Pf-5 pyoverdine match) | Present — NRPS + NI-siderophore region with high similarity to nunapeptin / nunamycin (Region 3.1); NRP-metallophore region reported (low Pf-5 pyoverdine match) | Both strains encode siderophore-type clusters; LMG 5496 includes a nunapeptin/nunamycin-like siderophore hit. |
| **Arylpolyene (APE)** | Present — Region 11.1 APE (low confidence; APE Ec match) | Present — Region 9.1 arylpolyene (low confidence) | APes (pigment / oxidative-stress protection) appear in both genomes (low similarity calls). |
| **NRPS-like / cryptic clusters** | Present — NRPS-like region(s) (e.g., Region 9.1 NRPS-like) and a few cryptic loci) | Present — Several NRPS-like / cryptic clusters (e.g., Region 8.1, 28.1 icosalide match) | LMG 5496 shows more numerous cryptic/NRPS-like loci with diverse MIBiG matches (icosalide, icosalide A/B, etc.). |
| **Small / other clusters (terpene, HCN, redox-cofactor, NAGGN, NAPAA)** | Present — terpene-precursor, hydrogen-cyanide regions, redox-cofactor, NAGGN, NAPAA appear across several contigs | Present — hydrogen-cyanide, terpene-precursor, NAGGN, NAPAA, redox-cofactor also detected in various regions | Both genomes contain several non-NRPS/PKS small clusters (HCN, terpenes, redox cofactors). |
| **Overall BGC count and diversity** | 15 regions (relaxed detection). Multiple NRPS, 1 trans-AT PKS, hybrid syringolin A, siderophore hits, APE, and several small clusters | 18 regions (relaxed detection). Multiple NRPS, several NRPS-like cryptic clusters, trans-AT PKS, syringolin A hybrid, nunapeptin/nunamycin match, APE, and several small clusters. | LMG 5496 reports slightly more regions (18 vs 15) and a higher number of distinct NRPS/NRPS-like hits; EC33 has many conserved hits but fewer total regions. |

### Comparative analysis of copper resistance genes

A presence/absence matrix of 48 copper resistance genes across 100 *Pseudomonas* strains, allows a direct comparison between our strains (LMG 5496 and EC33) and other reference and environmental isolates. Both *Pss* LMG 5496 and EC33 share the majority of their copper resistance machinery. They both possess 35 core genes (including *CopB*, *CusCBA*, various ATPases), which represent the basic chromosomal efflux and sequestration systems essential for surviving low-level environmental copper toxicity. Crucially, both strains are missing the 13 accessory genes (e.g., *pcoD*, *pcoC*, *CopP*), which are typically associated with high-level copper resistance acquired via mobile elements (plasmids or transposons) (see the heatmap presented in Figure 5). A specific difference exists in one copper-detoxifying enzyme variant: *Cu_oxidase_cueO_2* which is present in strain EC33, strain LMG 5496 lacks this specific version of the copper oxidase. Similarly, the non-pathogenic strain *P*. *lijiangensis* Phyllo_227 (often isolated from plant leaves or soil) also lacks this gene.

**Figure 5.**
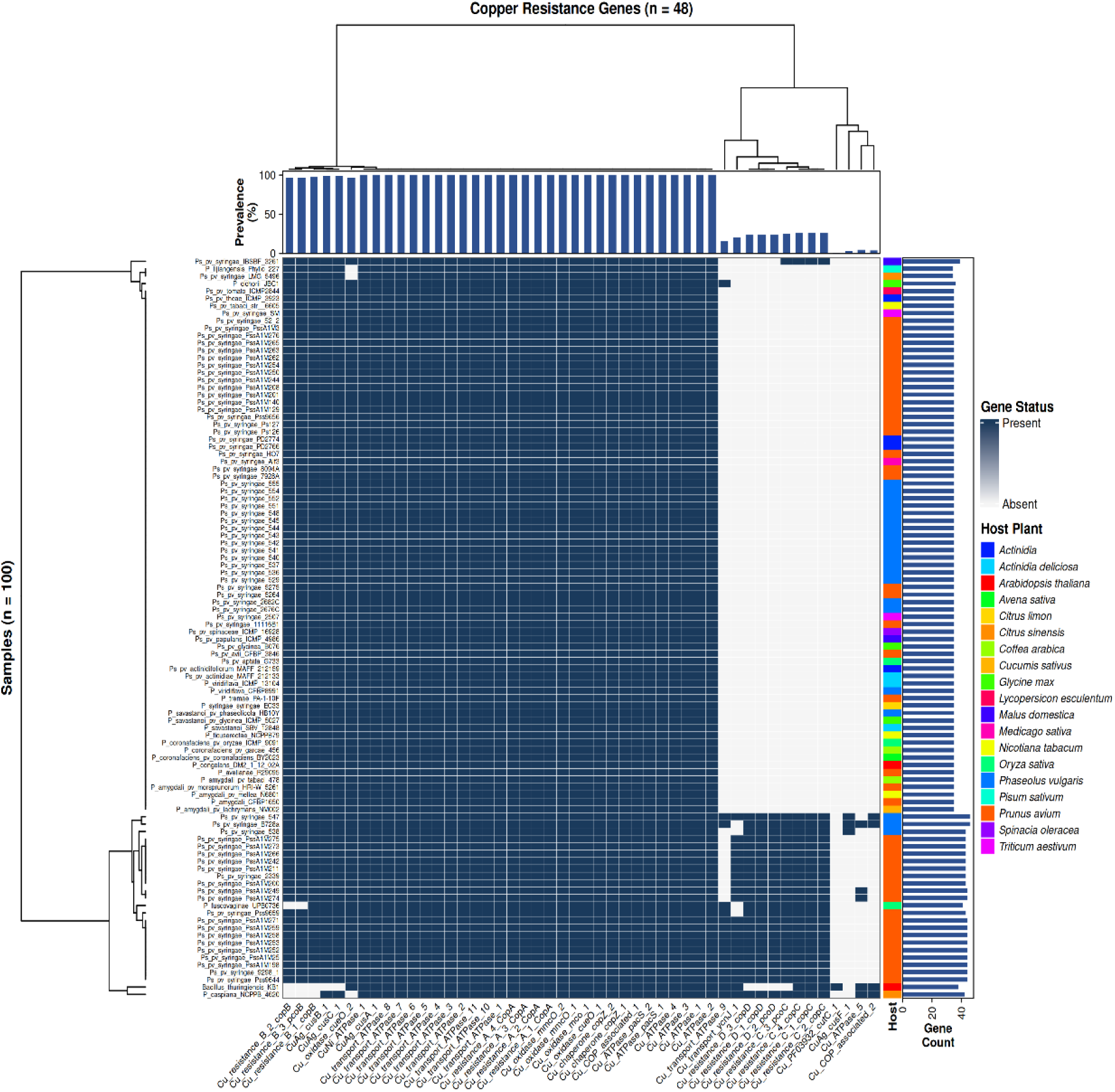
Presence/Absence Profiles and Hierarchical Clustering of Core and Accessory Copper Resistance Genes. This heatmap visualizes the presence or absence of 48 specific copper resistance genes across 100 samples.

### Prophage identification and features

Comparative prophage profiling revealed distinct prophage elements. PHASTEST detected two intact prophage regions in LMG 5496 and one in EC33, with no incomplete or questionable regions.

In LMG 5496, prophage Region 1 (35 kb; coordinates 599,780–634,836) resembled the sequence of *Pseudomonas* phage phi3 (NC_030940), while Region 2 (47.3 kb; 683,079–730,425) showed homology with the sequence of Enterobacteria phage Arya (NC_031048), suggesting recombination events. Both encode 32–39 proteins including integrase, capsid, tail, and lysis modules, confirming functional integrity.

EC33 harbored a single 38.4 kb intact prophage sequence (259,762–298,240), closely related to *Pseudomonas* phage YMC11/02/R656 (NC_028657), with 14 homologous proteins. All prophages exhibited GC contents of 57–59%, comparable to their host genomes (58.97%), consistent with stable integration and long-term co-evolution (Figure 6).

**Figure 6.**
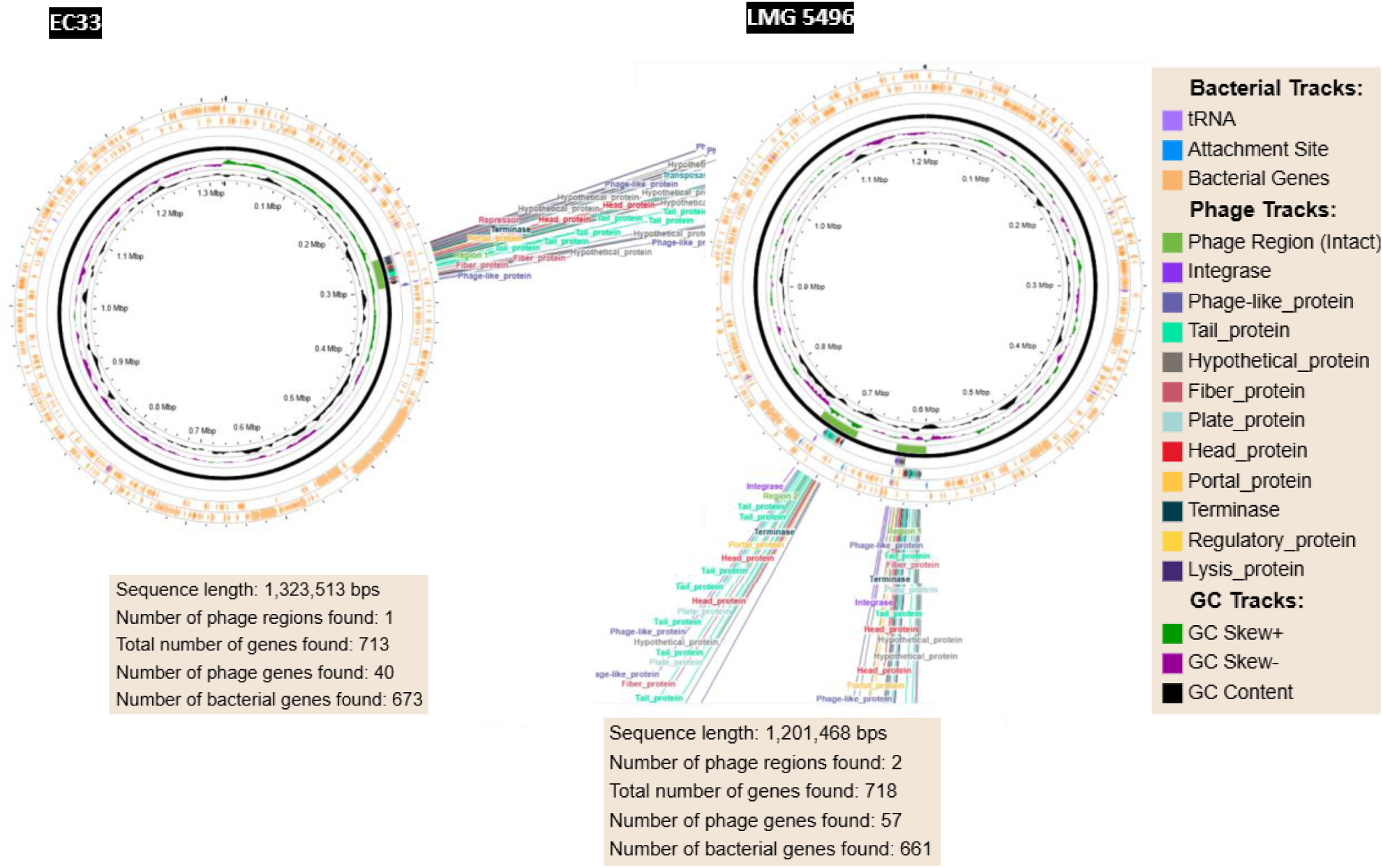
Comparative circular genome maps of *P*. *syringae* pv. *syringae* strains EC33 and LMG 5496 showing prophage regions. The circular diagrams represent the complete genomes of strains EC33 (left) and LMG 5496 (right). Prophage regions identified by PHASTEST are highlighted in green (intact), with phage-related genes annotated according to function. Sequence statistics are shown in the insets: total sequence length, number of prophage regions, total number of genes, and distribution of bacterial versus phage genes.

## Discussion

The high Average Nucleotide Identity (ANI) value ((≈ 100%) between strains EC33 and LMG 5496, alongside their extensive gene content homology, indicates a close relationship. The ANI analysis confirmed that strains LMG5496 and EC33 are closely related variants, exhibiting ≈ 100% sequence identity, indicating a stable core genome as witnessed over a period of 52 years However, divergence in their virulence gene repertoires suggests functional differentiation.

The identical T3E repertoires of EC33 and LMG 5496 indicate a conserved Type III effector arsenal. This shared set of effectors, including both canonical and accessory proteins, suggests that the two strains possess similar potential for host manipulation and virulence. The presence of conserved chaperones and structural T3SS components further supports the functionality and stability of the secretion system. The absence of strain-specific effectors implies that differences in pathogenicity or host range between these strains are unlikely to arise from variation in the high-confidence T3E repertoire.

Effector family annotations and analyses revealed both conserved and strain-specific effectors. In our dataset, heatmap clustering revealed distinct gene-level patterns associated with reduced virulence, loss of pathogenicity, or increased virulence, underscoring the genetic complexity and adaptive potential of *Pss* within the *P*. *syringae* species complex. PHI-base–guided comparative analysis of our strains showed a virulence architecture comprising: (i) a conserved Type III Secretion System (T3SS) core essential for infection competency, (ii) a variable effector module contributing to host adaptation, and (iii) a flexible regulatory network modulating virulence expression. In our strains, most effector gene disruptions were associated, according to PHI-base annotations, with reduced virulence or loss of pathogenicity, highlighting their central role in host colonization. A smaller subset of genes, including *BphP1* and *HrpV*, was linked to increased virulence, suggesting that regulatory fine-tuning may enhance aggressiveness under specific conditions.

These findings suggest that *Pss* combines a highly conserved core of virulence factors with plastic accessory modules to optimize infection strategies. The hierarchical effector distribution as established in earlier research on *Pseudomonas* (Swain et al., 2025) aligns with the “pan-effectorome” model, where core effectors provide basal pathogenic capability and variable/unique genes modulate host specificity and virulence intensity (Lindeberg et al., 2012). The co-occurrence in our dataset of conserved T3SS effectors with more flexible accessory genes, some likely horizontally acquired, supports a model of ongoing effector flux shaped by recombination and selective pressure from host immunity. This is consistent with previous findings showing recombination-driven diversification and effector turnover within *P*. *syringae* populations (Dillon et al., 2019). Together, our results reinforce the “pan-effectorome” concept by demonstrating that *Pss* maintains a stable virulence backbone while dynamically adjusting accessory modules to optimize infection under different ecological or host contexts.

Core effectors primarily correspond to conserved T3SS components (e.g. *hrp*, *hrc*, *avrE1*, *hopM1*), whose disruption consistently reduces virulence. Variable effectors, such as *hopZ1a*, *hopAO1*, and *avrRps4*, may mediate host-specific responses, while unique effectors located in the genomic islands suggest recent horizontal acquisition or strain-specific adaptation (Newfeld et al., 2025). The presence of an increased number of virulence-associated effectors in strain EC33 points to potential targets for future functional studies to elucidate their roles in pathogenicity. Additionally, identifying effectors linked to reduced virulence or loss of pathogenicity as is possibly the case for strain LMG 5496, may provide insights into the development of so-called laboratory strains or into host resistance mechanisms that can assist strategies for developing resistant plant varieties. Specifically, EC33 retained ten major virulence determinants, including the putative hypervirulence factor Q4ZSY4, while losing the avirulence gene Q52430. This combination likely contributes to an expanded host range and improved pathogenic fitness by preserving key pathogenicity functions while eliminating a host-recognition trigger.

Both strains possess genomic potential for secondary-metabolite production (siderophores, NRPS/PKS products, arylpolyenes), but LMG 5496 shows a higher number and diversity of NRPS/cryptic clusters, whereas EC33’s BGC set is slightly smaller and contains several highly conserved matches. This pattern suggests differences in biosynthetic repertoire that could affect iron acquisition, oxidative-stress tolerance and microbial competition.

The copper resistance gene analysis confirms that EC33 and LMG 5496 are fundamentally similar in their lack of acquired high-level resistance. Both *Pss* strains LMG 5496 and EC33 display a basal/chromosomal copper-resistance profile: they retain core chromosomal efflux and sequestration systems, but lack the accessory plasmid-borne operons commonly associated with high-level copper resistance (e.g., *pco*/*cop* mobile clusters). Chromosomal systems (where *cue*, *cus* and *cop*-like chromosomal genes are involved) provide intrinsic tolerance to environmental copper, while plasmid-borne *pco*/*cop* operons typically confer higher resistance and are often acquired by horizontal transfer in copper-exposed agro-ecosystems, due to the use of copper compounds as pesticide or growth promotor (Bondarczuk and Piotrowska-Seget, 2013; Lamichhane et al., 2018).

A single, robust genetic difference is the presence of a second multicopper oxidase variant (*Cu_oxidase_cueO_2*) in EC33, which is not present in LMG 5496. Multicopper oxidases such as *CueO* oxidize reduced copper species (Cu⁺) to less reactive Cu²⁺ and have been repeatedly shown to contribute to copper tolerance under aerobic conditions (Grass and Rensing, 2001). The presence of an extra *CueO* variant in EC33 plausibly improves detoxification capacity under pulse or chronic copper exposure and therefore represents a small but biologically meaningful advantage, even in the absence of plasmid-borne high-level resistance (Rowland et al., 2013). The core chromosomal machinery in both strains and an accessory detoxification enzyme only in EC33 fits a model of local selection and modular genome adaptation. EC33’s *CueO_2* may be a locally-selected accessory fine tuning resulting from strong and/or more persistent copper exposure in its environment (situation in Tunisia, 2014, see Abdellatif et al., 2020). Accessory genes like additional oxidases can appear by point divergence, gene duplication or horizontal acquisition from related taxa, and then be retained by selection where copper is used repeatedly. This mosaic acquisition of stress-response genes is distinct from the independent horizontal exchange of large virulence modules (e.g., T3SS effectors), and together these processes shape strain fitness in host and field contexts (Bondarczuk and Piotrowska-Seget, 2013).

Differences in prophage content between LMG 5496 and EC33 likely reflect microevolution driven by ecological context, host association, and/or geographic isolation. LMG 5496 harbored both *Pseudomonas*- and Enterobacteria-related intact prophages and prophage sequences, supporting a mosaic origin of mobile elements. Such cross-lineage phage associations are common in *Pseudomonas* spp., reflecting modular phage evolution and frequent HGT between bacteria sharing ecological niches (Koskella and Meaden, 2013; Touchon et al., 2017). Mosaic prophages may carry adaptive traits, including toxin production, surface structure modification, or resistance determinants, potentially contributing to host range expansion and virulence modulation (Hulin et al., 2023).

Despite fewer prophages, EC33’s larger genome suggests compensatory acquisition of non-phage genomic islands. The shared presence of intact prophages in both strains strongly indicates lysogenic competence. Such elements contribute to genomic plasticity, adaptive potential, and possible gene exchange relevant to host interaction, antibiotic resistance, stress tolerance, and environmental persistence (Varani et al., 2013; Shintani et al., 2022).

The presence of copper tolerance/resistance genes highlights the multifaceted adaptation strategies of *P*. *syringae*, combining efficient host colonization with tolerance to metal-induced oxidative stress. This dual adaptation may confer a selective advantage in agricultural settings where copper-based controls are prevalent (Lamichhane et al., 2018).

Our analysed genome sequences provide a foundation for studies on the molecular basis of adaptation to *Citrus* hosts. Understanding the balance of conserved core effectors, strain-specific accessory genes, prophages, and metal resistance determinants and their actual realization in the organism and environment may further illuminate the mechanisms underlying virulence, host specificity, and environmental persistence in *P*. *syringae* pv. *syringae*. Such insights can guide the development of improved disease management strategies, including breeding for resistance and searching alternatives for copper compounds.

## Acknowledgements

The sequencing of the bacterial strains was supported by the Laboratory of Bacteriology in the Flanders Research Institute for Agriculture, Fisheries and Food (ILVO), Plant Sciences Unit.

## Notes

https://gitlab.ilvo.be/stevebaeyen/pseudomonas_genomes_citrus_2025

### Competing Interest Statement

The authors have declared no competing interest.

